# Regulation of the plastochron by three *MANY-NODED DWARF* genes in barley

**DOI:** 10.1101/2020.12.15.422847

**Authors:** Ken-ichiro Hibara, Masayuki Miya, Sean Akira Benvenuto, Naoko Hibara-Matsuo, Manaki Mimura, Takanori Yoshikawa, Masaharu Suzuki, Makoto Kusaba, Shin Taketa, Jun-ichi Itoh

## Abstract

The plastochron, the time interval between the formation of two successive leaves, is an important determinant of plant architecture. We genetically and phenotypically investigated *many-noded dwarf* (*mnd*) mutants in barley. The *mnd* mutants exhibited a shortened plastochron and a decreased leaf blade length, and resembled previously reported *plastochron1* (*pla1*), *pla2*, and *pla3* mutants in rice. In addition, the maturation of *mnd* leaves was accelerated, similar to *pla* mutants in rice. Several barley *mnd* alleles were derived from three loci—*MND1, MND4*, and *MND8*. Although *MND4* coincided with a cytochrome P450 family gene that is a homolog of rice *PLA1*, we clarified that *MND1* and *MND8* encode an N-acetyltransferase-like protein and a MATE transporter-family protein, which are respectively orthologs of rice *GW6a* and maize *BIGE1* and unrelated to *PLA2* or *PLA3*. Expression analyses of the three *MND* genes revealed that *MND1* and *MND4* were expressed in limited regions of the shoot apical meristem and leaf primordia, but *MND8* did not exhibit a specific expression pattern around the shoot apex. In addition, the expression levels of the three genes were interdependent among the various mutant backgrounds. Genetic analyses using the double mutants *mnd4mnd8* and *mnd1mnd8* indicated that *MND1* and *MND4* regulate the plastochron independently of *MND8*, suggesting that the plastochron in barley is controlled by multiple genetic pathways involving *MND1, MND4*, and *MND8*. Correlation analysis between leaf number and leaf blade length indicated that both traits exhibited a strong negative association among different genetic backgrounds but not in the same genetic background. We propose that *MNDs* function in the regulation of the plastochron and leaf growth and revealed conserved and diverse aspects of plastochron regulation via comparative analysis of barley and rice.

**Author summary:** The number of leaves produced during a plant’s lifetime is major determinant of plant architecture and affects the efficiency of photosynthesis and crop productivity. The leaf number is dependent on the temporal pattern of leaf initiation at the shoot apical meristem, which is termed the plastochron. The genetic factors involved in plastochron regulation have been identified in several plant species. However, whether the functions of plastochron-related genes and their genetic pathways are universal or diversified among different plant species is unclear. In this study, we investigated *many-noded dwarf* (*mnd*) mutants in barley, which exhibited a shortened plastochron and a decreased leaf blade length. The mutant alleles used in this study were derived from three loci, *MND4, MND1*, and *MND8*, which encode a cytochrome P450 family protein, an N-acetyltransferase-like protein, and a MATE transporter-family protein, respectively. Phenotypic and expression analyses revealed that these three MND genes affect the leaf production rate and leaf maturation program, but their expression levels were interdependent. In addition, the plastochron and leaf growth are closely related but independently regulated. We also analyzed the expression patterns and knockout mutants of rice *MND* orthologs to clarify whether their biological functions are conserved in rice and barley. This study provides insight into the genetic mechanisms of plastochron control in grass species.

## Introduction

The spatiotemporal pattern of leaf initiation is a major contributor to the formation of plant architecture. The temporal pattern of leaf initiation is termed the plastochron; that is, the time interval between the initiation of two successive leaf primordia. The spatial pattern of leaf initiation along the shoot axis is referred to as phyllotaxy. Both patterns of leaf initiation are determined by the activity of the shoot apical meristem (SAM), which is the source of leaf primordia [1, 2]. Although the plastochron and the phyllotaxy are sometimes regulated by shared genetic components operating at the SAM, some genes are specific to one or the other of the patterns [1, 3].

*PLASTOCHRON1* (*PLA1*) specifically regulates the plastochron in rice (*Oryza sativa*) [4, 5]. Loss of function of *PLA1* causes rapidly emerging small leaves, resulting in more than twice the number of leaves compared to the wild type (WT). *PLA1* encodes a plant-specific cytochrome P450 family protein, CYP78A11 [5]. *Arabidopsis KLU* is also a CYP78A-family member and its loss-of-function mutant exhibits accelerated leaf initiation and produces small organs [6]. *PLA2* is another plastochron-regulating factor in rice [7]. *PLA2* encodes a MEI2-like RNA-binding protein, which is an ortholog of *TERMINAL EARl* (*TE1*) in maize (*Zea mays*) [8]. Although *TE1* plays a role in the regulation of phyllotaxy, accelerated leaf initiation in the loss-of-function mutant is shared between *te1* and *pla2*. *PLA3* has also been reported to regulate the plastochron. A loss-of-function mutant of *PLA3* exhibits not only a shortened plastochron but also pleiotropic phenotypes such as embryonic defects [9]. *PLA3* encodes a homolog of glutamate carboxypeptidase, which dissociates glutamate from small peptides [9]. *PLA3* is the rice ortholog of *Arabidopsis ALTERED MERISTEM PROGRAM1* (*AMP1*) [10] and maize *VIVIPAROUS8* [11]. Consequently, in rice, loss-of-function mutants of three *PLA* genes share the phenotypes of rapid leaf production, small leaf size, and aberrant inflorescence structure [4, 7, 9].

*SQUAMOSA PROMOTER BINDING PROTEIN-LIKE* (*SPL*) genes are plant-specific transcription factors, some of which are negatively regulated by *miR156* [12–15]. *SPL* genes are also involved in plastochron regulation. A double loss-of-function mutant of *AtSPL9* and *AtSPL15* in *Arabidopsis* exhibited a short plastochron [6, 16]. Conversely, the expression of an *miR156*-resistant form of *SPL9* caused a prolonged plastochron [6]. In grass species such as rice and maize, *SPL* genes and *miR156* have conserved functions in plastochron regulation. Accumulation of *OsSPL* transcripts caused a prolonged plastochron in rice and loss-of-function mutants of several *SPL* genes and plants overexpressing *miR156* exhibited a short plastochron in both rice and maize [17–23].

The *big embryo1* (*bige1*) mutant in maize exhibits an increased leaf and seminal root number in addition to a large embryo [24]. *BIGE1* encodes a MATE-type transporter that likely plays a role in the secretion of an unidentified small molecule in the trans-Golgi. *BIGE1* function is conserved between maize and *Arabidopsis*, because both single and double mutants of *BIGE1* homologs in *Arabidopsis* produced an increased number of leaves, and the introduction of *BIGE1-GFP* fusion genes into an *Arabidopsis bige1* mutant partially complemented the mutant phenotype [24].

Despite the identification of genes involved in the plastochron, the relationships among these genes and genetic pathways in plastochron regulation are unclear. In *Arabidopsis, KLUH/CYP78A5* and *AtSPL9/miR156* affect the plastochron and organ size in parallel genetic pathways [6]. In rice, a *pla1* and *pla2* double mutant exhibited enhanced phenotypes compared with each single mutant, suggesting that *PLA1* and *PLA2* function in independent pathways [7]. Although maize *BIGE1* is involved in the feedback regulation of a *CYP78A* pathway, the phenotypic effect of the interaction on the plastochron has not been elucidated [6]. In addition to the genetic pathways, knowledge of functional conservation of plastochron-related genes among plant species is fragmentary.

Barley (*Hordem vulgare*) is an important cereal crop for which considerable genetic resources are available, including collections of morphological and developmental mutants [25]. In addition, high-quality genome sequence information is available [26]. Thus, barley is an alternative model for grass molecular genetics with a less gene redundancy and is suitable for comparative studies with the other grass species. Here, we genetically and phenotypically characterized *many-noded dwarf* (*mnd*) mutants in barley, which were originally described in the 1920s as dwarf mutants with many nodes [27]. All *mnd* mutants exhibited a shortened plastochron, comparable to *pla* mutants in rice. We identified the genes responsible for *mnd* mutants and showed that the plastochron and leaf length are independently controlled by three genetic factors—*MND1, MND4*, and *MND8*. Here, we propose to assign a new gene symbol *mnd8* to a mutation that was found at a locus different from the seven previously described *mnd* loci in barley. We revealed that *MND8* encodes a MATE transporter-family protein, which is an ortholog of maize *BIGE1. MND1* encodes an N-acetyltransferase-like protein that reportedly regulates phase changes [28]. *MND4* had been reported as a cytochrome P450-family gene and a homolog of rice *PLA1* [29]. Our phenotypic and genetic analyses of the *mnd* mutants suggested functions for the three *MND* genes and the existence of complex genetic interactions among them. Furthermore, our comparative analysis of rice and barley clarified the diversity and conservation of plastochron genetic pathways.

## Results

### Plastochron and phyllochron of *mnd* mutants

The *mnd* mutants used in this study are listed in Table 1. Allelism tests revealed that these mutants carried alleles derived from at least three independent loci, *MND1, MND4*, and *MND8* (Table 1). *MND4* is also known as *HvMND* [29]. Among the mutants, three—OUM165, OUM169, and OUX051—which are presumed to contain the alleles *mnd8*_OUM165_, *mnd4*_OUM169_, and *mnd1*_OUX051_, respectively, and Akashinriki, an original cultivar of OUM165 and OUM169, were subjected to detailed genetic and phenotypic analyses.

**Table 1.** *mnd* mutants used in this study.

| Strain | Allele determined by allelism tests | Allele / mutation by sequencing | Mutation effect on protein | Background cultivar |
| --- | --- | --- | --- | --- |
| OUM165 | *mnd8* | *mnd8* / T→C | L→P | Akashinriki |
| OUM166 | n.d. | *mnd8* / G→A | G→N | ? |
| OUM168 | n.d. | *mnd8* / G→A | G→N | ? |
| OUM169 | *mnd4* | *mnd4* / G→A | G→D | Akashinriki |
| OUX051 | *mnd1* | *mnd1* / 8bp insertion | Truncated | Mesa |
| SM5 | *mnd1* | *mnd1* / 1bp deletion | Truncated | Kanto Nijo 29 |
| SM6 | *mnd1* | *mnd1* / 1bp deletion | Truncated | Kanto Nijo 29 |
| GSHO253 | *mnd1* | *mnd1* / 8bp insertion | Truncated | Mesa |
| GSHO2038 | n.d. | *mnd1* / 8bp insertion | Truncated | Bowman |
| GSHO2135 | n.d. | *mnd4* / G→A | R→K | Bowman |
| GSHO1798 | n.d. | *mnd4* / G→A | R→K | Akashinriki |
| NGB114540 | n.d. | *mnd1* / G→T | R→L | Kristina |
| NBG114547 | n.d. | *mnd8* / T→A | W→R | Bonus |
| NGB117205 | n.d. | *mnd8* / G→A | G→E | Bonus |

n.d. : not determined.

First, we evaluated the vegetative phenotypes. The three *mnd* mutants produced more leaves than Akashinriki at all stages of vegetative development (Fig 1A–1D). However, the extent of leaf emergence varied among the mutants and growth stages. In the WT and *mnd*s, leaf emergence was slow in the early vegetative phase and rapid in the late vegetative phase (Fig 1E). At 40 days after germination, the number of leaves that emerged in *mnd8*OUM165 and *mnd1*OUX051 was twofold that in the WT, and that in *mnd4*OUM169 was intermediate between those of the WT and other mutants, indicating that the three mutants have an increased leaf production rate (Fig 1E).

**Fig 1.**
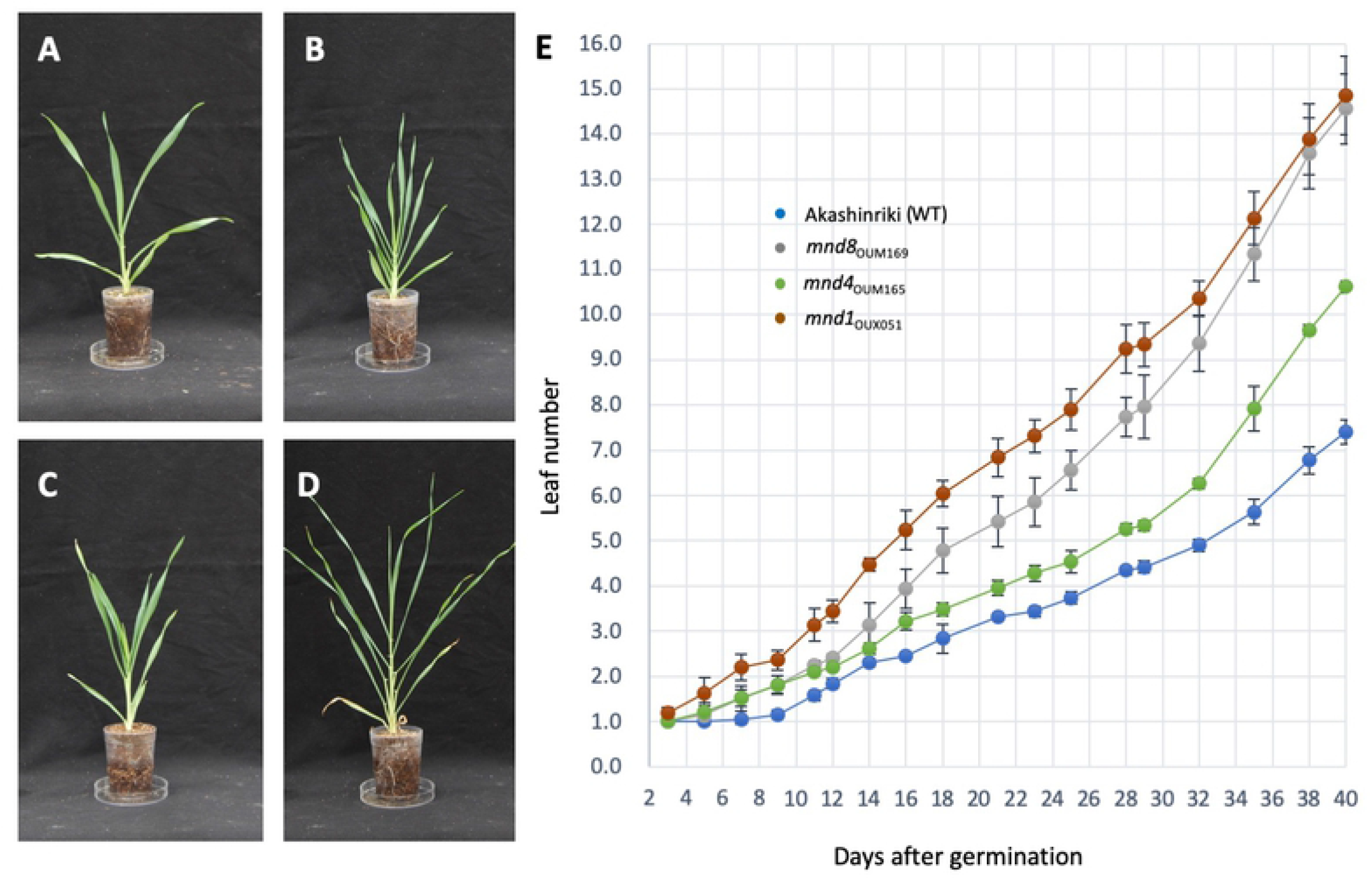
Vegetative phenotypes of *mnd* mutants. (A–D) Seedlings of the wild-type and *mnd* mutants at 1 month after germination. (A) Akashinriki, (B) *mnd8_OUM165_*, (C) *mnd4_OUM169_*, and (D) *mnd1_OUX051_*. (E) Changes in leaf number during vegetative development.

Two indices represent the temporal pattern of leaf production, the plastochron and the phyllochron [30]. The plastochron is the time interval between two successive leaves produced at the SAM, and the phyllochron is the time interval between two successive leaves emerging from the top of the former leaf sheath. Although the plastochron and phyllochron are equal in rice, the plastochron is shorter than the phyllochron in most other cereal crops [30]. To calculate the plastochron, we observed shoot apices of the WT and three *mnd* mutants at 1 and 2 weeks after germination (WAG) (S1 Fig). In most shoot apices of Akashinriki at 1 WAG, the P1 primordium was the fifth leaf protruding from the SAM (S1A Fig). At 2 WAG, the P1 primordium was the seventh leaf in most Akashinriki shoot apices (S1E Fig). Therefore, two new leaf primordia were produced from the SAM over 1 week in Akashinriki. In the *mnd* mutants, most of the P1 primordia were fifth leaves at 1 WAG, but eighth leaves at 2 WAG (S1B–1D, S1F–H Fig). The plastochrons of the three *mnd* mutants were shorter than that of the WT (Table 2). We also calculated the phyllochrons for the four growth periods. For the first period from 1 to 2 WAG, the phyllochron was longer than the plastochron in the WT and *mnd* mutants. Although the phyllochrons in the WT and *mnds* became shorter in the later periods, the difference between the WT and *mnd*s increased, suggesting that the leaf production rates of *mnd* mutants increased at later developmental stages. These results suggested that the three *mnd* mutants were defective in plastochron control, although the plastochron was shorter than the phyllochron in both the wild-type and *mnd* mutants.

**Table 2.**
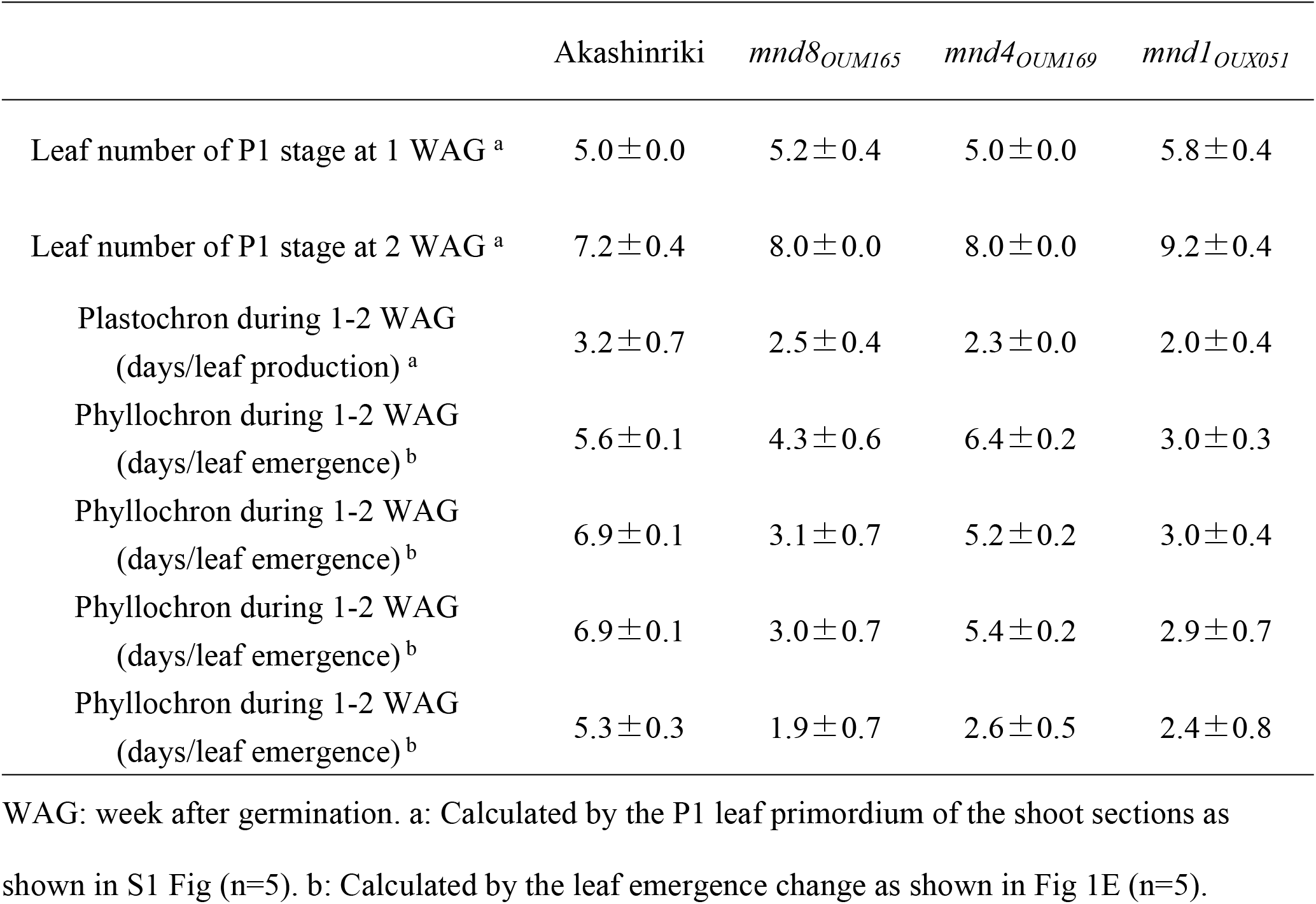
Characterization of leaf production and leaf emergence rate in *mnd* mutants.

### Growth and cell-division pattern of *mnd* leaf primordia

An increased leaf production rate may alter the spatial relationship among leaf primordia in the *mnds*. In other words, a rapid leaf production in the *mnds* may change the number of leaf primordia inside a leaf primordium at the same developmental stage. To assess this, we examined the shoot apices of the *mnd* mutants at 2 WAG under an electron microscope (S1 Fig and Fig 2). The relationships between successive leaf primordia in the *mnd*s were similar to that in the WT. Namely, when the P1 leaf primordium protruded from the flank of the SAM, the P4 leaf primordium gradually enclosed the inner leaf primordia in the WT (Fig 2A and 2B), and this relationship was conserved in the three *mnd* mutants (Fig 2C and 2F).

**Fig 2.**
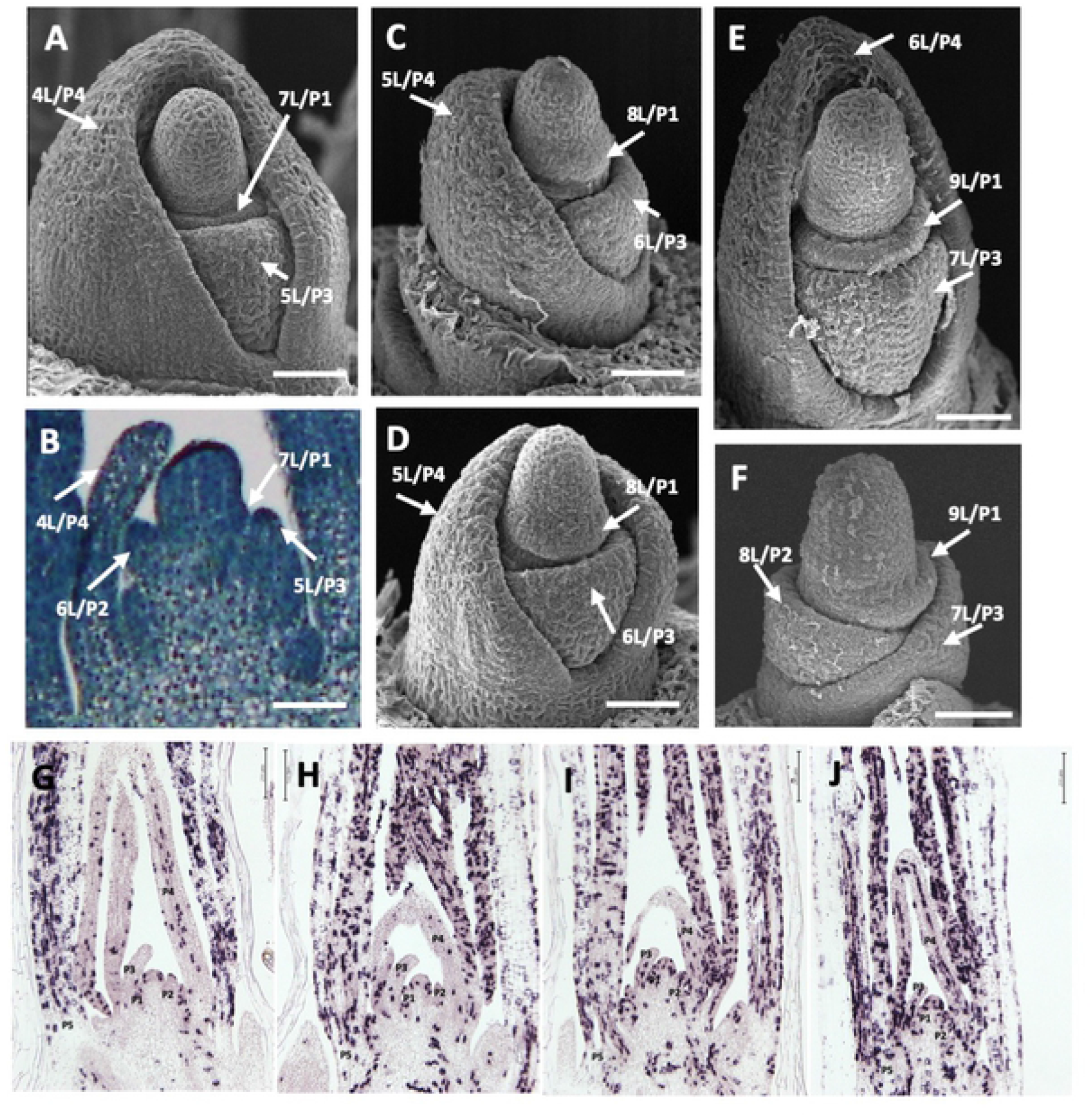
Spatial relationship between leaf-primordium stage and cell-division activity in *mnd* mutants. (A, C–F) Scanning electron micrographs of the shoot apex of *mnd* mutants at 2 weeks after germination. (A) Shoot apex of Akashinriki. (B) Longitudinal section of the shoot apex of Akashinriki corresponding to (A). Shoot apices in *mnd8_OUM165_* (C), *mnd4_OUM169_* (D), and *mnd1_OUX051_* (E). (F) Shoot apex of *mnd1_OUX051_* from which the 6L/P4 primordium was removed. xL indicates the x^th^ leaf and Px is the order of leaf emergence from the shoot apical meristem. (G–J) Expression pattern of histone *H4* in the shoot apex of wild-type and *mnd* mutants at 10 days after germination according to *in situ* hybridization. (G) Akashinriki, (H) *mnd8_OUM165_*, (I) *mnd4_OUM169_*, and (J) *mnd1_OUX051_*. Bars: 50 μm in A–F and 200 μm in G–J.

To understand the growth pattern of leaf primordia in the *mnds*, cell division around the shoot apex was investigated by *in situ* hybridization using the cell-division biomarker histone *H4* (Fig 2G–2J). The number of histone *H4* signals around the shoot apex in the *mnd* mutants was generally higher compared with the WT. In particular, histone *H4* signals in the *mnd1* were remarkably increased (Fig 2G and 2J). However, patterns of density of histone *H4* signals in the *mnd4* and *mnd8* mutants were similar to that in the WT; namely, cell division activity was relatively low at P1 to P4 and enhanced in P5-stage leaf primordia in the WT, *mnd4*, and *mnd8*. Therefore, stage-specific developmental events in leaf primordia in the *mnd4* and *mnd8* are likely unchanged, although leaf stage progressions could be more accelerated in the *mnd1*. Thus, the barley *mnds* exhibited not only rapid leaf production but also rapid leaf development, as do *pla* mutants in rice [7].

### Molecular identification of three *MND* genes

The three *mnd* mutants exhibited similar phenotypes, and the affected traits were comparable to those of *pla* mutants in rice. Accordingly, the three *MND* genes in barley could be counterparts of the three rice *PLA* genes [5, 7, 9].

To identify the three *MND* loci, we first determined the nucleotide sequences of *MND4/HvMND* (HOR5Hr1G081060) in OUM169, a homolog of rice *PLA1* [29] (S2A Fig). We detected a single nucleotide polymorphism between Akashinriki and *mnd4*OUM169. The mutation was a G-to-A single base change (G425A) causing a Gly-to-Asp amino-acid substitution in the first exon of *MND4* (Fig 3A, Table 1). This amino acid is conserved among angiosperms, suggesting it to be responsible for the phenotype of *mnd4*_OUM169_ (S3 Fig). We detected an identical mutation in the other two *mnd* mutants, *mnd4*_GSHO2135_ and *mnd4*_GSHO1798_, that caused an Arg-to-Lys amino-acid substitution, which has been reported previously [29] (Fig 3A, Table 1, S3 Fig). Because we found multiple *mnd4* alleles linked to a shortened plastochron phenotype, *MND4/HvMND* regulates the plastochron in barley.

**Fig 3.**
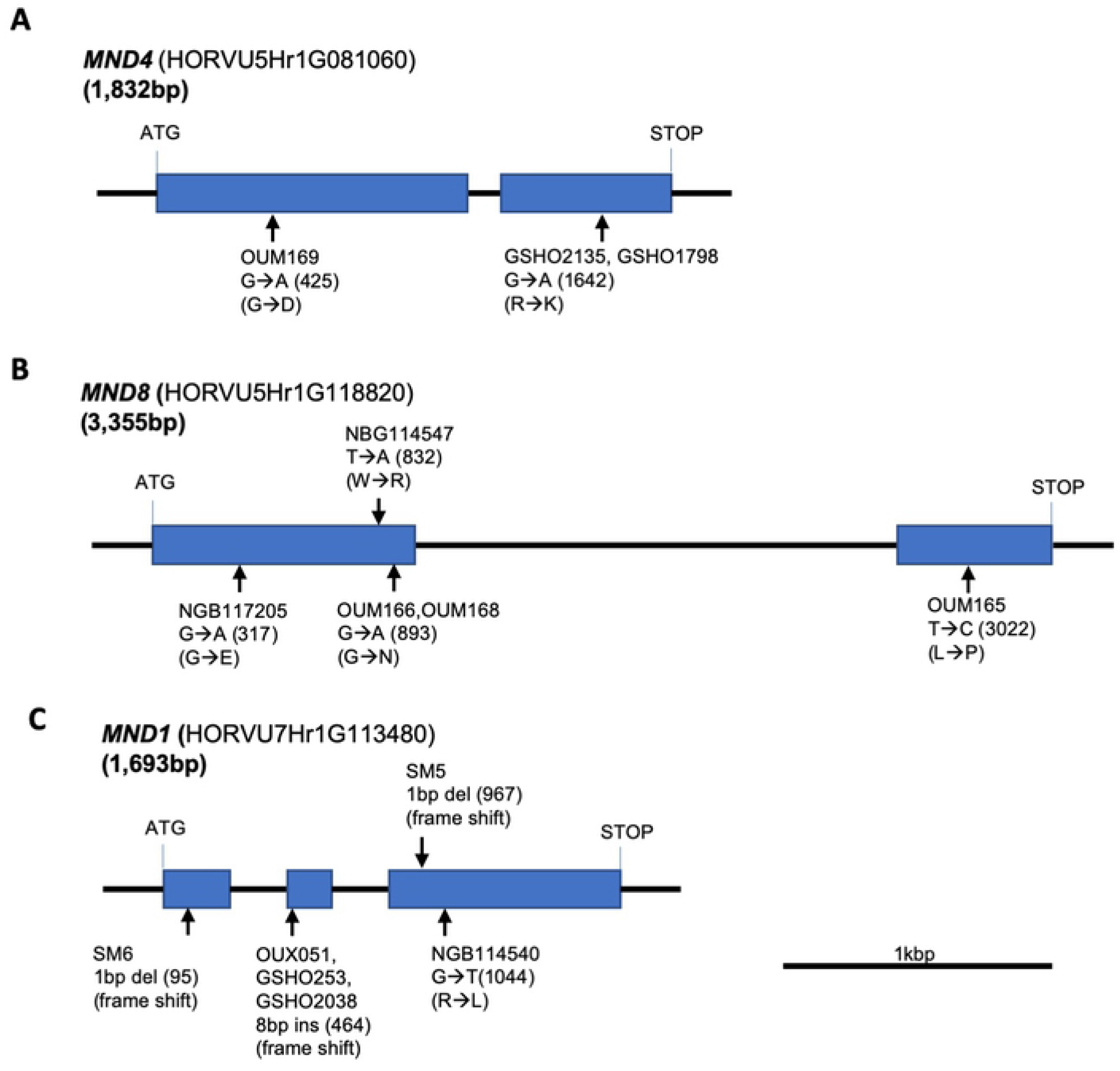
Genomic structure of MND genes. (A–C) Genomic structure and mutation points of MNDs. (A) *MND4*, (B) *MND8*, and (C) *MND1*. Boxes indicate exons, arrows indicate mutation points.

Next, to identify candidates for *MND1* and *MND8*, we tested the barley ortholog of *PLA2: HvPLA2* (HORVU3Hr1G091930). Rice *pla2* exhibits a phenotype similar to those of barley *mnd* mutants [7]. We determined the nucleotide sequences of *HvPLA2* in various *mnd* mutant and WT pairs. A database search showed that *HvPLA2* is located in barley chromosome 3H, which is syntenic to rice chromosome 1 carrying *PLA2*, but our mapping showed that *MND1* and *MND8* are located on chromosomes 7H and 5H, respectively. We found no mutation causing an amino-acid change or frameshift in *HvPLA2* in *mnd8*_OUM165_ and *mnd1*_OUX051_, suggesting that neither *MND1* nor *MND8* is *HvPLA2* (S6 Fig).

A barley ortholog of rice *pla3* was next investigated because of its similar phenotype [9]. The barley ortholog of *PLA3* (HORVU5Hr1G103900), *HvPLA3* in rice chromosome 3, in located on syntenic chromosome 5H. We mapped *MND8* to the distal region of the long arm of barley chromosome 5H, but this position deviated distally from the *HvPLA3* candidate interval. No mutation was detected in the nucleotide sequence of *HvPLA3* in *mnd8*_OUM165_. Sequencing of *HvPLA3* in *mnd1*OUX051 revealed two point mutations causing amino-acid substitutions, and a six-base deletion causing two amino-acid deletions in *HvPLA3* in *mnd1*_OUX051_. Although one of the point mutations and the deletion mutation did not change the amino acids conserved among angiosperms, a C-to-A point mutation (C773A) in the first exon caused a Pro-to-Gln amino-acid substitution (P258Q) at a conserved position (S6 and S7 Figs). However, we did not find nucleotide polymorphisms between Akashinriki and the other *mnd* mutants with *mnd1* alleles, i.e., *mnd1*_GSHO253_ and *mnd1*_SM6_. Based on these inconsistent mapping and sequencing results, *HvPLA3* is not responsible for *mnd1*.

Because it is unlikely that *MND1* and *MND8* are *HvPLA2* or *HvPLA3*, we searched for other candidate genes that regulate the plastochron based on reports in species other than rice. Rough mapping in barley showed that *mnd8* is located 2 cM distal to the barley EST marker k08652 (syn. HORVU5Hr1G112990), whose map position is 253.1 cM in the distal region of the long arm of chromosome 5H according to the EST map [31]. Around the mapped region, we found a candidate gene, a homolog of *BIGE1*, which regulates not only embryo size but also the leaf initiation rate in maize [24]. We subsequently examined the nucleotide sequence of the *BIGE1* homolog (HORVU5Hr1G103900), which encodes a MATE transporter-family protein. We found a T-to-C single base change (T3022C) mutation in the second exon of the gene in *mnd8*_OUM165_ that caused a Leu-to-Pro amino-acid substitution (L404P) (Fig 5B, Table 1, S4 Fig). The nucleotide sequence of the gene in four *mnd* mutants—OUM166, OUM168, NBG114547, and NGB117205—revealed three independent mutations causing amino-acid substitutions (G106E, W278R, and G298N) in the coding region. Accordingly, we concluded that this MATE transporter*-*like gene is *MND8*. A phylogenetic analysis indicated that *MND8* is an ortholog of maize *BIGE1* (S2 Fig).

Next, we mapped *mnd1* using publicly available simple sequence repeat (SSR) markers in barley [32]. Barley *mnd1* was located in the distal region of the long arm of barley chromosome 7H, flanked by the SSR markers GBM1419 and Bmac156. Using this map information, we searched for *mnd1* candidate genes. We identified as a strong candidate for *MND1* (HORVU7Hr1G113480) a homolog of *HOOKLESS1* (*HLS1*), which encodes an N-acetyltransferase-like protein. Although there are no reports on *HLS1* homologs regulating the plastochron in *Arabidopsis*, multiple *Arabidopsis* mutants of *HLS1* homologs exhibited excess leaf production and a small leaf size [33–35]. One *HLS1* homolog in barley was located near the distal arm region of chromosome 7HL, where *mnd1* was mapped. Sequence analysis of this gene revealed that *mnd1*_OUX051_ carries an eight-nucleotide deletion in the second exon starting at position 464, causing a frameshift and introducing a premature stop codon (Fig 3C, Table 1, S5 Fig). Sequencing of four *mnd* mutants with the *mnd1* alleles (*mnd1*_SM5_, *mnd1*_SM6_, *mnd1*_GSHO253_, and *mnd1_GSHO2038_*) and one other *mnd* mutant, NGB114540, revealed an eight-base insertion, two independent one-base deletions, and one amino-acid substitution in the coding region (Fig 3C, Table 1, S5 Fig). In addition, among the mutations in *MND1*, the eight-base insertion in *mnd1*_OUX051_, *mnd1*_GSHO253_, and *mnd1*_GSHO2038_ was identical to that reported in *mnd1.a* [28]. Therefore, we concluded that this *HLS1*-like gene is *MND1*. A phylogenetic analysis indicated that *MND1* is an ortholog not of *HLS1* but of *GW6a*, a quantitative trait locus that regulates grain weight in rice [36] (S2 Fig).

Accordingly, the three *MND* genes—*MND4, MND8*, and *MND1—*encode CYP78A family, MATE transporter-family, and N-acetyltransferase-like proteins, respectively.

### Expression pattern of the three *MND* genes

To evaluate their regulation of the plastochron and leaf growth, the expression patterns of the three *MND* genes around the shoot apex were investigated using *in situ* hybridization (Fig 4). *MND4* expression was observed both in the SAM and leaf primordia (Fig 4A). The expression pattern in the SAM was complex, with expression detected in small patches of the SAM flank. One patch was observed at the boundary between the SAM and P1 leaf primordia. Strong *MND4* expression was observed in the basal part of the adaxial and abaxial sides of P1–P4 leaf primordia, including the boundary between successive leaf primordia (Fig 4A). Expression of *MND8* was weak in the SAM and younger leaf primordia, and was not observed in the shoot apex (Fig 4B).

**Fig 4.**
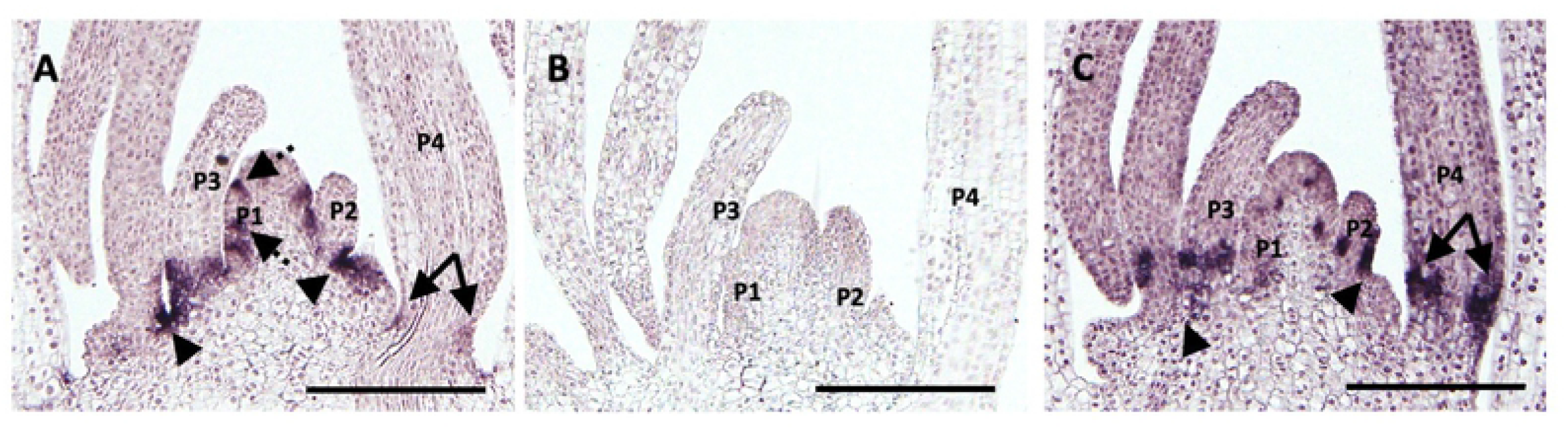
Expression pattern of the three *MND* genes in the shoot apex at 10 days after germination. (A) *MND4*, (B) *MND8*, and (C) *MND1*. The P1–P4 leaf primordia are labeled. Arrowheads indicate gene expression at the boundaries between leaf primordia. The dashed arrow in (A) indicates expression at the boundary between the shoot apical meristem and P1 leaf primordium. Two-headed arrows indicate expression at the leaf base. Note the relative position of expression at the leaf base differs between (A) and (C). Bars: 200 μm.

**Fig 5.**
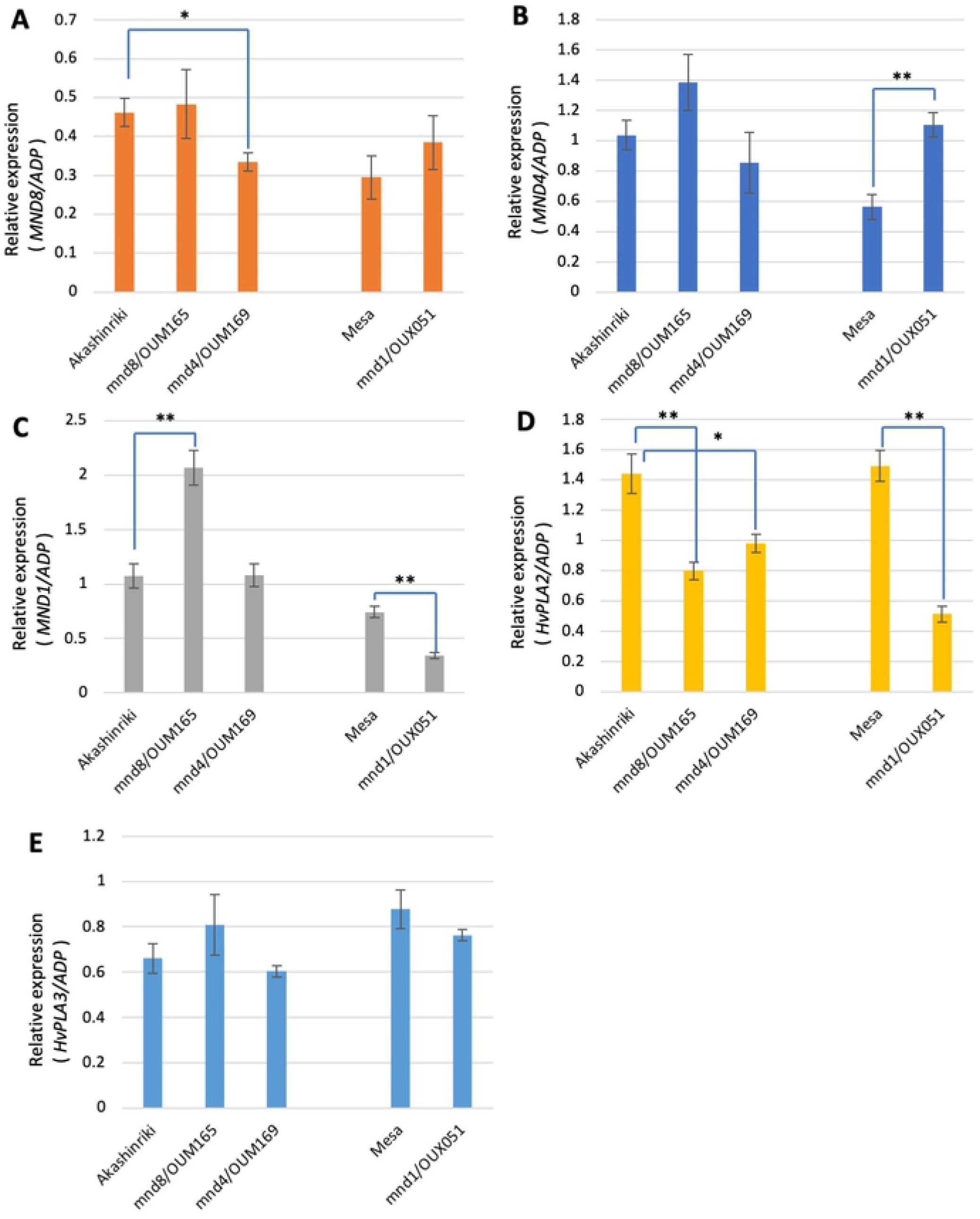
Expression change in *MND* and *HvPLA* in *mnd* mutant backgrounds. Relative expression levels of *MND8* (A), *MND4* (B), *MND1* (C), *HvPLA2* (D), and *HvPLA3* (E). *ADP-ribosylation factor 1-like protein* (*ADP*) was used as the internal control. Double and single asterisks indicate a statistically significant difference compared to the wild type (WT; *t*-test, P < 0.01 and 0.05, respectively).

The expression pattern of *MND1* was similar to that of *MND4;* that is, expression was observed in small patches of the SAM and the basal part of leaf primordia (Fig 4C). However, the area of expression was not identical to that of *MND4*. First, *MND1* expression was detected not at the boundary between the SAM and P1 leaf primordia but in the inner region of the SAM. Second, gene expression at the boundary between successive leaf primordia was marked for *MND4*, but not for *MND1*. Finally, *MND1* expression in the basal parts of leaf primordia was more apically shifted compared to *MND4*.

These observations suggest that *MND4* and *MND1* regulate the plastochron and leaf growth by means of their expression in limited but distinct regions of the SAM and leaf primordia.

### Effect of *mnd* mutations on the expression level of the three *MND* genes

To examine whether *MND* genes regulate other *MND* genes, we compared the expression levels of *MND* genes among three *mnd* mutant backgrounds and WT Akashinriki and Mesa, with the former the original cultivar of *mnd4*_OUM169_ and *mnd8*_OUM165_ and the latter that of *mnd1*_OUX051_ (Fig 5). Real-time PCR revealed that the expression of *MND8* was slightly decreased in *mnd4*_OUM169_ (Fig 5A). This indicates that *MND4* positively regulates *MND8* expression. The expression level of *MND4* was slightly upregulated in *mnd8*_OUM165_ and significantly upregulated in *mnd1*_OUX051_, suggesting that *MND1* negatively regulates *MND4* expression (Fig 5B). The expression level of *MND1* was upregulated in *mnd8*_OUM165_ and downregulated in *mnd1*_OUX051_, implying that *MND8* negatively regulates *MND1* expression and that *mnd1*_OUX051_ affects the accumulation of *MND1* mRNA (Fig 5C). In addition, we examined the expression of *HvPLA2* and *HvPLA3* (Fig 5D and 5E). Although *HvPLA3* expression was not significantly altered in the three *mnd* mutants (Fig 5E), *HvPLA2* expression was downregulated in the *mnd* mutants (Fig 5D). This suggests that the three *MND* genes positively regulate *HvPLA2* expression.

Accordingly, the expression levels of the three *MND* genes in addition to that of *HvPLA2* were affected by functional defects in other *MND* genes.

### Genetic interactions among *MND* genes

To examine the genetic interactions among *MND* genes, we generated double mutants between *mnd8*_OUM165_ and *mnd4*_OUM169_ as well as *mnd8*_OUM165_ and *mnd1*_OUX051_ (Fig 6A and 6B). We measured the leaf number and leaf blade length in the F_3_ population at 2 months after germination and determined the genotypes (Fig 6C and 6D). Among the segregants of the F_3_ population of *mnd8*_OUM165_ and *mnd4*_OUM169_, double homozygotic plants for *mnd8*_OUM165_ and *mnd4*_OUM169_ produced the largest number of leaves among the nine genotypes (Fig 6A and 6C). Although the difference in leaf number between single *mnd8*_OUM165_ homozygotic plants and double homozygotic plants was slight, mature double-mutant plants exhibited an enhanced phenotype relative to the *mnd8*_OUM165_ single mutant (S8 Fig), indicating that the *mnd8*_OUM165_ allele has an additive effect with *mnd4*_OUM169_. Among the segregants of the F_3_ population produced from *mnd8*_OUM165_ and *mnd1*_OUX051_, an additive effect between the two alleles on leaf number was evident; that is, the *mnd8*_OUM165_ *mnd1*_OUX051_ double mutant had the largest number of leaves among the genotypes (Fig 6B and 6D).

**Fig 6.**
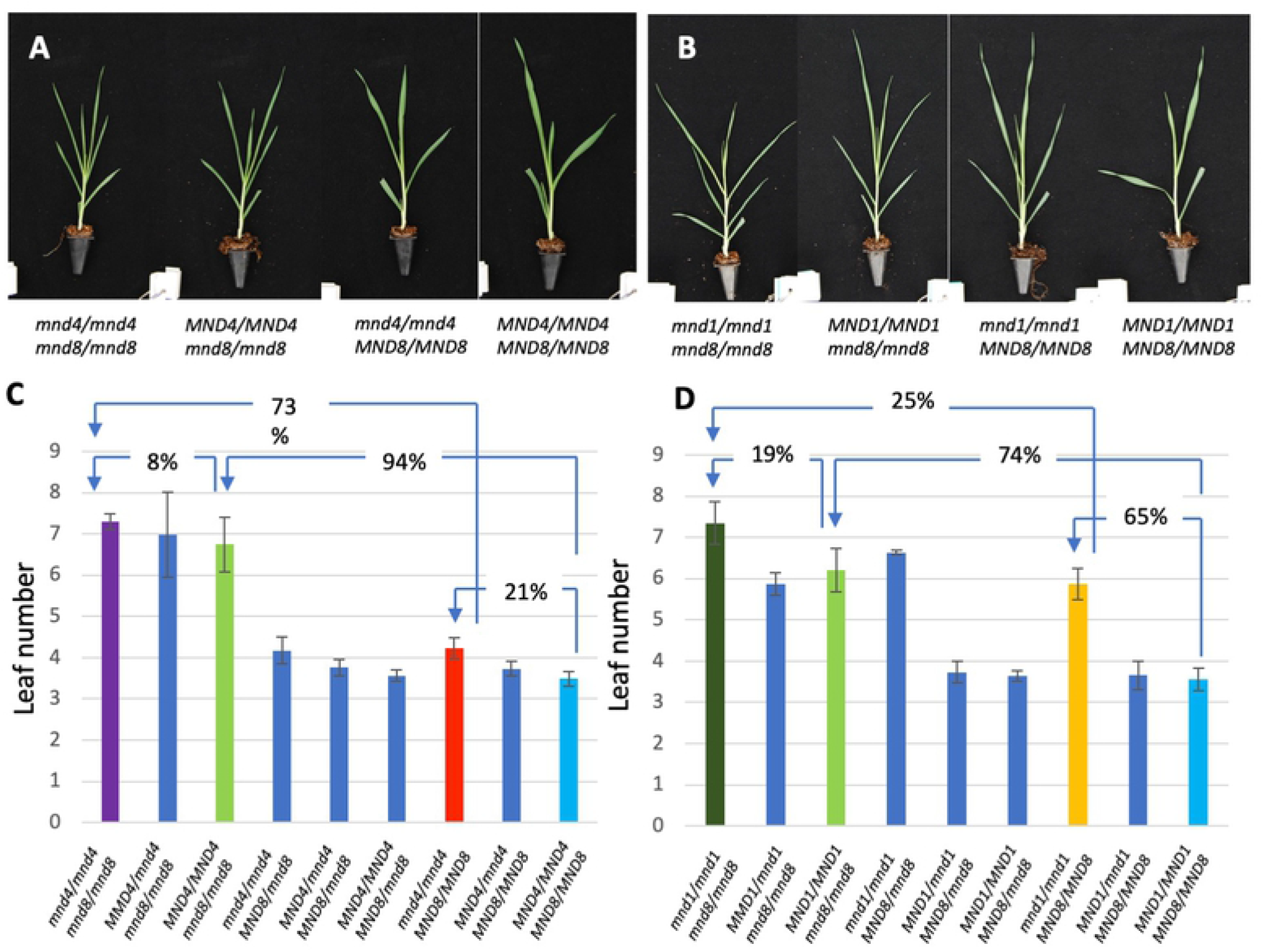
Effect of *mnd* double mutations on leaf number. (A, B) Seedling phenotypes of single and double mutants of *mnd4*_OUM169_ and *mnd1*_OUX051_ (A) and *mnd4*_OUM169_ and *mnd8*_OUM165_ (B) at 2 months after germination. Genotypes are indicated below. (C, D) Effect of *mnd4*_OUM169_ and *mnd1*_OUX051_ (C) as well as *mnd4* _OUM169_ and mnd8_OUM165_ (D) alleles on leaf number. Arrows and percentages indicate the effect of homozygotic mutations of *mnds* on the rate of increase in leaf number.

The *mnd* alleles had different effects on the increase in leaf number depending on their genotype (Fig 6C and 6D). For example, *mnd8*_OUM165_ increased leaf numbers in the WT and *mnd4*_OUM169_ backgrounds by 94% and 73%, respectively. Similarly, *mnd4*_OUM169_ increased leaf numbers in the WT and *mnd8*_OUM165_ backgrounds by 21% and 8%, respectively (Fig 6C). The same tendency was observed for combinations of *mnd8*_OUM165_ and *mnd1*_OUX051_ (Fig 6D). Accordingly, the effect of a single *mnd* allele on the increase in leaf number was diminished by the accumulation of other *mnd* alleles.

Therefore, *MND8* regulates the plastochron independently of *MND4* and *MND1*. In addition, there are genetic or developmental mechanisms that moderately affect the leaf production rate in the short plastochron background.

### Relationship between the plastochron and leaf-blade length in the *mnd* mutant

The shortened plastochron in most plastochron-related mutants is reportedly accompanied by the production of small leaves [4, 6, 7, 24]. However, the correlation between leaf production rate and leaf length is unknown. Therefore, the leaf number at 2 months after germination and the length of the second leaf blade of F_3_ segregants produced from *mnd8*_OUM165_ and *mnd4*_OUM169_ and from *mnd8*_OUM165_ and *mnd1*_OUX051_ were measured (Fig 8). A strong negative correlation between leaf number and leaf-blade length was observed in all F_3_ segregants for both *mnd8*_OUM165_ and *mnd4*_OUM169_ crosses and *mnd8*_OUM165_ and *mnd1*_OUX051_ crosses (Fig 7A and 7B). This correlation was caused by the strength of the allelic effect on leaf production (Fig 7C and 7D). By contrast, no negative correlations were detected among groups of segregants with identical genotypes (S9 Fig). In addition, among various WT and *mnd* mutant strains, no correlation between leaf number and the maximum length of the leaf blade was observed (S10 Fig, S1 Table).

**Fig 7.**
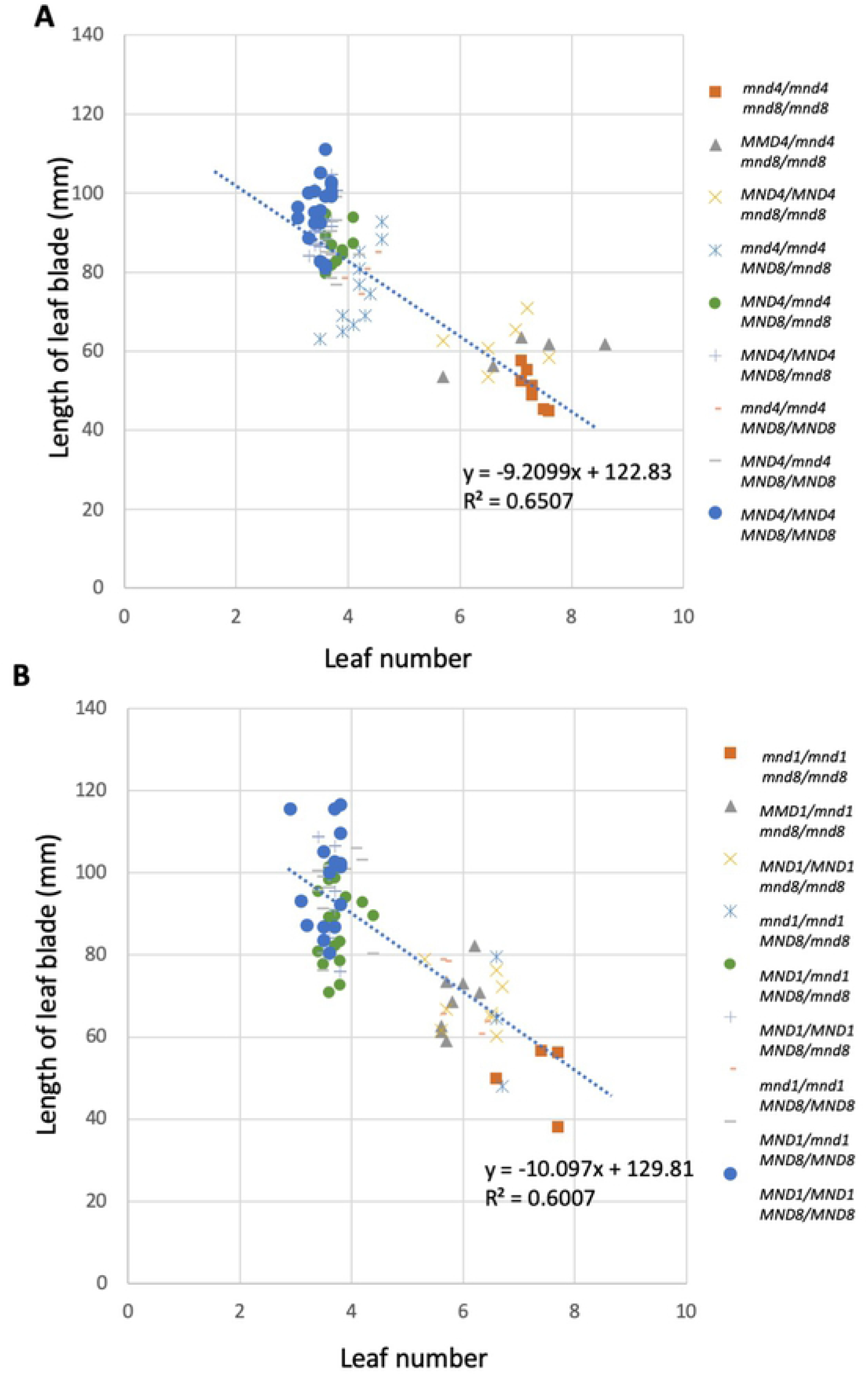
Relationship between leaf number and leaf-blade length. (A, B) Scatter plots of leaf number at 2 months after germination and the length of the second leaf blade in segregated plants from *mnd4 _OUM169_* × *mnd8_OUM165_* (A) and *mnd1_OUX051_* × *mnd8_OUM165_* (B) crossings. The linear regression line and coefficient of determination (R^2^) are indicated.

**Fig 8.**
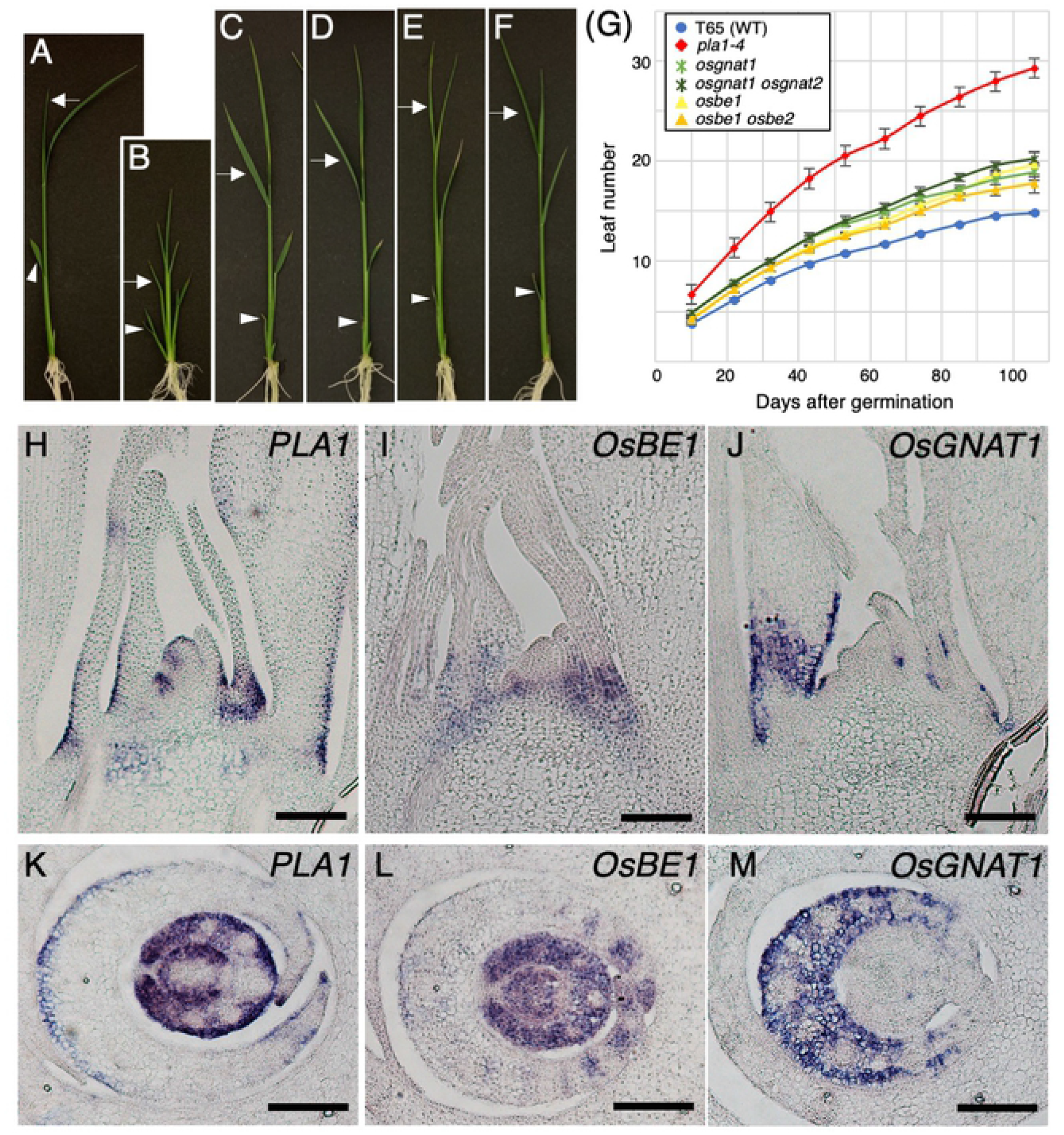
*MND* orthologs in rice. (A–F) Seedlings of the WT and rice mutants in *mnd* orthologs at 10 days after germination. (A) T-65, (B)*pla1-4*, (C) *osbe1*, (D) *osbe1 osbe2*, (E) *osgnat1*, and (F) *osgnat1 osgnat2*. Arrowheads and arrows indicate the second and fourth leaf blades, respectively. (G) Changes in leaf number during vegetative development. (H–M) Localization of *PLA1* (H, K), *OsBE1* (I, L), and *OsGNAT1* (J, M) mRNAs in longitudinal (H–J) and cross (K–M) sections of the shoot apex of 10-day-old seedlings. Bars: 100 μm.

These observations indicate that the plastochron and leaf-blade length are positively regulated by *MND* genes, but they are controlled independently rather than by alterations in the plastochron or leaf length.

### Effect of rice *MND* orthologs on plastochron regulation

Our results demonstrated that *MND8* and *MND1* regulate the plastochron in barley, but whether the rice homologs of these genes are involved in plastochron regulation is unknown. Therefore, we constructed knockout mutants of rice *MND8* and *MND1* orthologs using CRISPR/Cas9 (S11 Fig.). A phylogenetic analysis indicated that two orthologs of barley *MND8* and *MND1* are present in rice (S2 Fig; two rice orthologs, *OsBE1*/Os03g0839200 and *OsBE2*/Os12g0552600, for barley *MND8* and two rice orthologs, *OsGNAT1/Os06g0650300* and *OsGNAT2*/Os02g0180400, for barley *MND1*). We generated single and double mutants of the rice orthologs and calculated the plastochron (Fig 8A–8G). The number of leaves increased in *osgant1, osbe1, osgnat1 osgnat2*, and *osbe1 osbe2* compared to the WT, but to a lesser extent compared to *pla1-4* (Fig 8G, S12 Fig). This indicates that rice *OsBE1* and *OsGNAT1* are involved in plastochron regulation, as are their orthologs in barley, but their contribution to the plastochron is of lesser magnitude than that of *MND* in barley.

*In situ* hybridization showed that *OsBE1* and *OsGNAT1* were expressed around SAMs, particularly at the base of young leaf primordia (Fig 8H–8M), whereas *OsBE2* and *OsGNAT2* expression was not observed (data not shown). We next compared the expression patterns of *PLA1, OsBE1*, and *OsGNAT1. PLA1* and *OsBE1* expression was similar; *PLA1*, but not *OsBE1*, was expressed at the base of the P1-related region in the SAM (Fig 8H and 8I). In addition, *OsBE1*, but not *PLA1*, expression was detected in the inner tissue of P3 and the stem tissue (Fig 8K and 8L). *OsGNAT1* expression was strongest at the base of P3 and the boundaries of leaf primordia (Fig 8J and 8M). Expression of the three genes partially overlapped, but none was expressed in the vascular bundle (Fig 8K–8M). By contrast, there were differences in the expression patterns of the rice and barley orthologs, although *MND4* and *PLA1* expression was similar. Punctate expression of *MND1* was observed in the SAM (Fig 4C), whereas *OsGNAT1* expression was detected in the P3 leaf primordium but not in the SAM (Fig 8J and 8M). *MND8* did not exhibit a specific expression pattern (Fig 4B), but *OsBE1* was expressed around the shoot apex (Fig 8I and 8L).

These results indicate functional conservation between the barley and rice orthologous genes (*MND8* versus *OsBE1* and *MND1* versus *OsGNAT1*).

## Discussion

### Three *MND* genes regulate the plastochron via similar developmental pathways, but have unrelated functions and genetic pathways

Our results indicate that the many-noded phenotype of *mnd* mutants results from the rapid production of leaf primordia, which is caused by loss-of function of three independent loci, *MND4*, *MND8*, and *MND1*. Although the magnitude of the plastochron reduction differed among the mutant alleles, the overall phenotypes of all *mnd* mutants during vegetative development were similar. In addition, the panicle-development phenotypes were comparable. In the normal panicle of barley, a non-branched single axis of the panicle is produced, but diverged and multiple panicles were observed in *mnd8_OUM165_* and *mnd1_OUX051_* (S11 Fig). A panicle abnormality was also recognized in *mnd4*_OUM169_, which exhibited the weakest vegetative-development phenotype among the *mnd* mutants. Therefore, common abnormalities were exhibited during vegetative and reproductive development, indicating that the three *MND* genes have similar roles in barley development.

Despite their similar biological functions, the three *MND* genes encode unrelated proteins. *MND4* encodes a CYP78A monooxygenase whose substrates are unknown [29]. Although CYP78A genes are important for development in several species [29, 37–41], the synthetic or metabolic pathways in which CYP78A is involved are unknown. *MND1* encodes an N-acetyltransferase-like protein and is the closest homolog of *GW6a* in rice [36]. Because *GW6a* exhibits histone H4 acetyltransferase activity, *MND1* also likely regulates the transcription of downstream genes by acetylating histone H4. Furthermore, MND8 is an ortholog of maize BIGE1, a transporter implicated in the secretion of an unidentified small molecule [24]. Accordingly, it is assumed that MND4, MND1, and MND8 are involved in the synthesis or metabolism of unknown factors, transcriptional regulation of downstream genes, and transportation of unidentified molecules, respectively. Although a close relationship between the CYP78A pathway and *BIGE1* has been proposed [24], there is no direct evidence that *BIGE1* is associated with the transportation of CYP78A-related molecules so far. In fact, although we did not investigate the relationship between *MND4* and *MND1*, our genetic analysis indicates that *MND4* and *MND1* regulate the plastochron independently of *MND8*. Accordingly, at least two different genetic pathways regulate the plastochron in barley. This is also the case for plastochron regulation by three *PLA* genes in rice. The *pla1, pla2*, and *pla3* mutants have a short plastochron and a small leaf size, and conversion of the primary rachis into a shoot occurs in all three, but it has been proposed that *PLA1, PLA2*, and *PLA3* regulate the plastochron independently [7, 9]. On the other hand, *Arabidopsis AMP1* and *CYP78A5/7*, which are orthologs of *PLA3* and *PLA1*, respectively, act on a common downstream process [42].

Therefore, the developmental program controlling leaf initiation, leaf growth, and panicle development in rice and barley is regulated by multiple and independent genetic pathways involving *PLA* and *MND* genes.

### Interactions of *MND* genes

At least two independent genetic pathways regulate the plastochron, and complex genetic interactions were suggested by the expression analysis. In short, *MND8* negatively regulates *MND1*, *MND1* negatively regulates *MND4*, and *MND4* positively regulates *MND8*. It was also noted that *MND4* expression was slightly upregulated, but not significantly, in *mnd8*. Such a regulatory relationship between CYP78A and a *MND8*-related transporter has also been reported in maize; *i*.*e*., the expression of some CYP78A family genes was upregulated in the *bige1* mutant background [24]. In maize, *BIGE1* is required for feedback regulation of CYP78A family genes. If this is also the case in barley, the upregulation of *MND4* and *MND1* might have been the result of a deficiency in feedback regulation caused by the loss of *MND8* function. By contrast, overexpression of rice *GW6a*, an ortholog of *MND1*, and the *mnd1.a* mutation affected the expression levels of thousands of genes in rice and barley, respectively [28, 36]. Because *GW6a* is predicted to be a transcriptional regulator that modulates chromatin status, a change in *MND4* expression in *mnd1* could be a direct or indirect effect of loss of histone acetyltransferase activity by MND1. Although MND4 and MND8 do not directly affect gene transcription, they modulate that of other genes by influencing downstream events. In addition to the three *MND* genes, expression of *HvPLA2* was downregulated in the three *mnd* mutants. This indicates that *HvPLA2* is involved in plastochron regulation downstream of the three *MND* genes.

Phenotypic analysis of double mutants revealed that the effect of *MND* genes on plastochron regulation was dependent on the number of *mnd* mutant alleles present. Although it is unknown whether this phenomenon is a result of genetic interactions among the *MND* genes or a developmental restriction of plastochron regulation at the SAM, complex genetic and developmental interactions are implicated in the regulation of the plastochron.

### *MND* genes function in leaf production and growth

In shortened-plastochron mutants, rapid leaf production is accompanied by small leaves [4, 6, 7, 24]. In *pla1* and *pla2* mutants in rice, a model of the effect of leaf production on leaf size was proposed. According to the model, rapid leaf production by *pla* mutants is not the result of the loss of *PLA* function in the SAM, but an indirect effect of accelerated leaf maturation in leaf primordia [7]. The model was based on the observation that *PLA* genes were expressed in leaf primordia but not in the SAM, although the mechanism by which the change in leaf-primordia maturation affects leaf production is unknown.

Our barley analysis supports the above notion. The relative developmental stage and cell-division activity of successive leaf primordia were maintained or increased in the *mnd* mutants. Therefore, leaf production and leaf-primordia maturation were accelerated, as for *pla* mutants in rice. Accordingly, *MND* genes maintain the leaf-maturation schedule. However, if the leaf production rate is affected only by leaf maturation, as suggested by the rice model, the correlation between leaf number and leaf-blade length should be solid. However, we showed that leaf number and leaf-blade length were not always correlated in plants of identical genotype, implying that the plastochron and leaf size are differentially regulated. In fact, expression of *MND4* and *MND1* was observed both in the SAM and leaf primordia. *MND* expression in the SAM may be required for the suppression of leaf initiation independently of expression in the leaf primordia, which is necessary for the suppression of leaf maturation. This is supported by the fact that enhanced expression of the maize *PLA1* ortholog in leaf primordia affected leaf size but not leaf number [41], whereas overexpression of *PLA1* via the introduction of an increased gene copies resulted in a prolonged plastochron and an increased leaf size [43]. Furthermore, maize *PLA1* genes control the duration of cell division in the leaf primordia [41], which is consistent with the rapid leaf maturation observed in the *mnd4* mutant in this study.

The mechanisms by which *MND* genes regulate leaf initiation in the SAM and leaf growth in the leaf primordia are unknown. Auxin may be associated with control of the duration of cell-division activity by maize PLA1 [41]. It is also probable that auxin triggers leaf initiation via *MND*-mediated regulation, because the local auxin concentration is important for leaf initiation in the SAM [1, 44]. It is possible that the limited *MND1* and*MND4* expression domain in the SAM is important for the auxin flow or concentration required for proper leaf initiation. Analysis of auxin dynamics in the mutant SAM may provide insight into the role of auxin in the temporal regulation of leaf initiation by *MNDs*.

### Conservation and diversity of genetic pathways in rice and barley

Although several genetic factors regulating the plastochron have been identified in rice, maize, *Arabidopsis*, and other species, our understanding of their functional conservation among plant species is inadequate. We characterized *mnd* mutants in barley and identified the responsible genes. It was somewhat surprising that two of the three *MND* genes were not homologs of three *PLA* genes in rice. At least five genes are involved in developmental pathways underpinning plastochron regulation in grasses. Moreover, three of the five orthologs perform similar functions in rice and barley. However, there was a considerable difference in the magnitude of their effects between the two species. For example, loss-of-function of *MND1* and *MND8* markedly shortened the plastochron in barley, but that of the rice orthologs *OsGNAT1* and *OsBE1* resulted in a slight effect, even for double loss-of-function of closely related paralogs. This might have been caused by differences in genetic redundancy and functional diversification of gene families between the two species. Namely, other homologs in different clades compensate for the function of plastochron regulation in rice. In fact, the functional ortholog of *MND4* in rice is *PLA1*, but it is not the closest homolog among the CYP78A members [29]. It is predicted that *Os09g0528700* is a phylogenetic ortholog of *MND4* in rice, which is specifically expressed in roots and the embryo (http://ricexpro.dna.affrc.go.jp), unlike *PLA1*. In terms of the functions of *MND8-related* MATE transporter-family genes, *BIGE1* in maize regulates both the leaf initiation rate and embryo size [24]. However, *mnd8* in barley did not exhibit a phenotype change in terms of embryo size (S14 Fig). In addition, *OsBE1*, a rice ortholog of *MND8*, exhibited specific expression around the shoot apex, unlike *MND8*. Accordingly, the phenotypic effect and expression pattern of the closest homologs of *MND* genes differ among grass species.

Our understanding about a conserved function of plastochron-related genes among grass species is still insufficient. Identifications of *HvPLA2* and *HvPLA3* mutants in barley and comparative analyses with their rice counterparts would advance an understanding of the conservation and diversity of plastochron-related genes.

## Materials and Methods

### Plant materials and growth conditions

The barley *mnd* mutant and WT strains used in this study were listed in Table 1. Mutant and wild-type seeds were grown in soil under natural conditions, and the plants were sampled and their phenotypes evaluated at predetermined timepoints. For the double mutant analysis, the genotypes of the F3 plants were identified using PCR-based genotyping.

The *pla1-4* mutant and WT rice plants were grown on soil or MS medium containing 3% sucrose and 1% agar at 28°C under continuous light. Transgenic plants were grown in a biohazard greenhouse with temperatures of 30°C in the daytime and 25°C at night.

### Histological and morphological analysis

Samples of the mutant and WT plants were fixed with FAA (formaldehyde:glacial acetic acid:50% ethanol, 2:1:17) for 24 h at 4°C for histological analysis, or with PFA (4% [w/v] paraformaldehyde and 1% Triton X in 0.1 M sodium phosphate buffer) for 48 h at 4°C for *in situ* hybridization. They were then dehydrated in a graded ethanol series, after which the ethanol was substituted with 1-butanol, and the samples were embedded in Paraplast^®^ Plus (McCormick Scientific). The samples were sectioned at a thickness of 8 μm using a rotary microtome. The sections were stained with hematoxylin for histological analysis. After staining, the sections were mounted with Poly-Mount^®^ (Polysciences, Inc.) and observed under a light microscope.

For scanning electron microscopy, plant materials were fixed in PFA for 24 h and dehydrated in an ascending ethanol series, which was then gradually replaced with 3-methyl-butyl-acetate. Samples were critical-point dried, sputter-coated with platinum, and observed under a scanning electron microscope (S-4800, Hitachi) at an accelerating voltage of 10 kV.

### *In situ* hybridization

Paraffin sections were prepared as described above. For digoxigenin-labeled antisense RNA probes, those for histone *H4 (HvH4:* HORVU5Hr1G086620), *MND4, MND8*, and *MND1* in barley and *OsBE1* and *OsGNAT1* in rice were prepared using cDNAs and specific primers (S2 Table). The *PLA1* probes were prepared as described previously [5]. *In situ* hybridization and immunological detection using alkaline phosphatase were performed according to the methods of Kouchi and Hata [45].

### Identification of *MND* genes and *PLA2* and *PLA3* orthologs in barley

The nucleotide sequences of *MND4, MND8, MND1, HvPLA2*, and *HvPLA3* in the *mnds* and WT were determined using sequence information from the IPK Barley Blast Server (https://webblast.ipk-gatersleben.de/barley_ibsc/) and Phytozome (https://phytozome.jgi.doe.gov/).

For phylogenic analyses, the amino-acid sequences of MND4, MND8, and MND1 homologs in various plant species were obtained from the Phytozome database. The amino-acid alignment was carried out using GENETYX software (Genetyx), and the phylogenic tree was constructed based on the neighbor-joining method with 1,000-replicate bootstrapping using Molecular Evolutionary Genetics Analysis (MEGA X; [46]).

### Quantitative real-time PCR

RNA was extracted from the shoot apices of barley seedlings at 2 WAG using TRIzol^®^ reagent (Invitrogen). The extracted RNA was treated with Recombinant DNase I (TaKaRa), and cDNA was synthesized using the High-Capacity cDNA Reverse Transcription Kit (Life Technologies). Quantitative real-time PCR was performed with the StepOne^™^ Real-Time PCR System (Life Technologies) using TaqMan Fast Universal PCR Master Mix and FAM-labeled TaqMan probes for each gene. *ADP-ribosylation factor 1-like protein* (*ADP*) was used as the internal standard [47]. In all experiments, we analyzed three technical and three biological replicates. The primers and TaqMan probes for *MND1, MND4, MND8, ADP, HvPLA2*, and *HvPLA* are listed in S3 Table.

### Generation of knockout alleles of rice *MND* orthologs using the CRISPR/Cas9 system

The CRISPR/Cas9 system was used to generate knockout alleles for OsBE1, OsBE2, OsGNAT1, and OsGNAT2, which are orthologs of *MND1* and *MND8* in rice. The target sites were selected using the CRISPR-P program (http://cbi.hzau.edu.cn/crispr/) [48] (S11 Fig). The single-guide RNA (sgRNA) cloning vector (pZK_gRNA) and all-in-one binary vector (pZH_OsU6gRNA_MMCas9) harboring sgRNA, Cas9, and NPTII were provided by Masaki Endo [49]. The pZH_OsU6gRNA_MMCas9 vector including target-guide RNA for MND orthologs was constructed as described previously [49].

The constructs were introduced into *Agrobacterium tumefaciens* strain EHA105 and transformed into cultivar Taichung-65 (T-65) calli via *Agrobacterium-mediated* transformation. Mutations and transgenes in each transformant were confirmed by sequencing and PCR-based detection (S11 Fig).

## Data availability

The GenBank accession numbers for the genes in the text are *HvPLA2*, LC593235; *HvPLA3*, LC593236; *MND4*, LC593237; *MND1*, LC593238; *MND8*, LC593239; *OsBE1*, LC593240; *OsBE2*, LC593241; *OsGNAT1*, LC593242; and *OsGNAT2*, LC593243.

## Acknowledgments

The barley resources were provided by the National Bioresource Project-Barley, Okayama University, Japan; USDA, Aberdeen, Idaho; Nordic Genetic Resource Center, Sweden; and Tochigi Agricultural Experiment Station, Japan. This work was supported in part by the Ministry of Education, Culture, Sports, Science and Technology (MEXT) as part of a Joint Research Program implemented at the Institute of Plant Science and Resources, Okayama University, Japan.

## Supporting Information

**S1 Fig. Changes in the leaf primordia of *mnd* mutants at 1–2 weeks after germination.** Inner structure of the wild-type and *mnd* mutants at 1 week (A–D) and 2 weeks (E–H) after germination. (A, E) Akashinriki, (B, F) *mnd8_OUM165_*, (C, G) *mnd4_OUM169_*, and (D, H) *mnd1_OUX051_*. xL indicates the x^th^ leaf and Px indicates the order of leaf emergence from the shoot apical meristem. col, coleoptile. Bars: 200 μm

**S2 Fig. Phylogenetic tree of MND proteins.** Phylogenetic tree of MND proteins from several angiosperms. (A) MND4, (B) MND8, and (C) MND1. Numbers above the branches are bootstrap values from 1,000 replicates. At, *Arabidopsis thaliana;* Zm, *Zea mays;* Os, *Oryza sativa;* Hv, *Hordeum vulgare*. Red and blue underlining indicates MNDs in barley and orthologs in rice, respectively.

**S3 Fig. Amino-acid alignment of MND4 and its homologs.** Alignment of MND4 and its homologous proteins in several angiosperms used in S2 Fig. The effect of each *mnd* mutation is indicated in red. Black and gray, 100% and more than 50% identical amino acids, respectively. At, *Arabidopsis thaliana*; Zm, *Zea mays*; Os, *Oryza sativa*; Hv, *Hordeum vulgare*

**S4 Fig. Amino-acid alignment of MND8 and its homologs.** Alignment of MND8 and its homologous proteins in several angiosperms used in S2 Fig. The effect of each *mnd* mutation is indicated in red. Black and gray, 100% and more than 50% identical amino acids, respectively. At, *Arabidopsis thaliana*; Zm, *Zea mays*; Os, *Oryza sativa*; Hv, *Hordeum vulgare*.

**S5 Fig. Amino-acid alignment of MND1 and its homologs.** Alignment of MND1 and its homologs in the angiosperms used in S2 Fig. The effect of each *mnd* mutation is indicated in red. Black and gray, 100% and more than 50% identical amino acids, respectively. At, *Arabidopsis thaliana;* Zm, *Zea mays;* Os, *Oryza sativa;* Hv, *Hordeum vulgare*.

**S6 Fig. Genomic structure of *HvPLA2* and *HvPLA3*.** Boxes indicate exons. DNA polymorphisms between Akashinriki and OUX051 in the *HvPLA3* genomic structure are indicated by arrows.

**S7 Fig. Amino-acid alignment of HvPLA3 and its homologs.** Alignment of HvPLA3 and its homologs in several angiosperms. The effects of mutations in OUX051 are indicated in red. Black and gray, 100% and more than 50% identical amino acids, respectively. LOC, *Oryza sativa;* GRMZM, *Zea mays;* Solyc, *Solanum lycopersicum;* Medtr, *Medicago truncatula;* AT, *Arabidopsis thaliana*.

**S8 Fig. Mature plant phenotypes of *mnd* double mutants.** (A) Segregated plants of the F_3_ population derived from the F_2_ seeds of *mnd4 _OUM169_* × *mnd8_OUM165_* crossings. (B) Segregated plants of the F_3_ population derived from the F_2_ seeds of *mnd1_OUX051_* × *mnd8_OUM165_* crossings. The genotypes of the plants are indicated.

**S9 Fig. Relationship between leaf number and leaf-blade length among plants of the same genotype.** (A, B) Scatter plots of leaf number at 2 months after germination and the length of the second leaf blade with the same genotype from *mnd4 _OUM169_* × *mnd8_OUM165_* (A), and *mnd1_OUX051_* × *mnd8_OUM165_* (B) crossings. The linear regression line and the coefficient of determination (R^2^) are indicated.

**S10 Fig. Relationship between leaf number and maximum leaf-blade length.** Scatter plot of leaf number at 100 days after germination and the maximum length of the leaf blade. Average values of the traits of five plants were used. The linear regression line and coefficient of determination (R^2^) are indicated.

**S11 Fig. CRISPR/Cas9-mediated mutagenesis of rice *MND* orthologs.** (A–D) Genomic structures and target sites of rice MND orthologs. (A) *OsGNAT1*, (B) *OsGNAT2*, (C) *OsBE1*, and (D) *OsBE2*. Boxes indicate exons. Arrows indicate target sites (protospacer adjacent motif [PAM] sequences [red characters] and guide sequences [blue characters]) and directions starting from each PAM sequence. The lower sequence is that of the mutant used in the experiment. Bars: 500 bp.

**S12 Fig. Leaf number of wild-type (T-65) plants and CRISPR/Cas9-induced mutants at 10 days after sowing.** Data are presented as the means ± SDs (n ≥ 5).

**S13 Fig. Panicle phenotypes of *mnd* mutants.** Arrows indicate elongated bracts or ectopic shoot-like structures; arrowheads indicate branched panicles.

**S14 Fig. Seed phenotypes of *mnd* mutants.** (A) Akashinriki, (B) *mnd4 _OUM169_*, (C) *mnd8_OUM165_*, and (D) *mnd1OUX051*. Bars: 2 mm.

**S1 Table. Leaf number and leaf size in the wild type and *mnd* mutants.**

**S2 Table. Primers used to produce *in situ* hybridization probes.**

**S3 Table. Primers and TaqMan probes used for real-time PCR.**

